# Temporal compression of gonadal growth preserves reproductive timing and soma-to-germline scaling under stress

**DOI:** 10.64898/2026.10.14.759048

**Authors:** Peter Lenart, Ioana Gheorghe, Sacha Psalmon, Benjamin Towbin

## Abstract

How organisms maintain timely reproduction under stress is central to life history plasticity and remains an unmet challenge in biomedicine. Life history theory predicts that organisms preserve fitness across environments by reallocating resources between somatic and reproductive growth^1–6^, yet direct tests of this prediction have been limited by the lack of longitudinal life history data. Here, using *Caenorhabditis elegans*, we combine longitudinal video microscopy of gonadal growth, mathematical modelling, and genetic perturbations to show that reproductive timing is preserved through temporally dynamic control of reproductive organ growth. Under stress, the onset of gonad growth was delayed early in development but later accelerated relative to somatic growth, thereby preserving reproductive-to-somatic size ratios at adulthood. Modelling suggests this delay-and-acceleration strategy to minimize the time to maturity under stress. Consistently, disrupting compensatory gonadal growth acceleration, either through genetic perturbation of the JNK/p38 stress signalling pathway or by uncoupling of reproductive and somatic development, reduced gonad size and delayed reproduction. Together, our results show that reproductive robustness under stress arises from dynamic regulation of reproductive growth rather than a fixed shift in resource allocation, defining a previously unrecognized mechanistic principle for environmentally plastic sexual maturation.

## Introduction

Effective reproduction requires the precise coordination of somatic growth and reproductive organ development across changing environments^7,8^. Although the timely maturation of reproductive organs is critical for fertility, premature investments in reproductive growth impose energetic costs during juvenile stages, when these organs provide no reproductive benefit^9,10^. How developing organisms balance this tradeoff between somatic and reproductive growth to achieve timely reproduction remains poorly understood.

In many species, including humans, the timing and progression of reproductive maturation are strongly influenced by environmental conditions. Following dietary changes in modern societies, the age of puberty has declined substantially in recent decades ^11–13^, whereas childhood malnutrition continues to be associated with delayed sexual maturation^14–20^. However, different pubertal milestones are often differentially affected by these environmental conditions^12,21^. For example, the onset of breast development (Tanner B2 stage), an early pubertal milestone, has advanced more than the later milestone of menarche in recent decades ^21^ and is more strongly associated with the body-mass-index (BMI) ^22,23^. The mechanism underlying these non-linear shifts in sexual maturation dynamics are not understood.

A prominent explanation for such plasticity in reproductive development is that plasticity evolved under selective pressure to maintain reproductive fitness under variable environmental conditions ^24–26^. Consistent with this view, theoretical models of life history plasticity predict that plastic allocation of resource between somatic and reproductive tissues can optimize selective fitness ^1–6,27^. However, despite the central role of these ideas in organismal and evolutionary biology, direct tests relating fitness-relevant traits to resource allocation remain limited, in part because these processes are difficult to measure and manipulate *in vivo*.

Relating developmental responses to long-term reproductive success requires longitudinal and non-invasive measurements, which are not feasible in most systems. Here, we address this challenge by using long-term live imaging in *Caenorhabditis elegans* to quantify somatic and gonadal growth in individual animals. We identify a dynamic developmental strategy in which gonadal growth is initially slow but later accelerates sharply. Environmental stress delays the onset of gonad expansion while enhancing its rate relative to somatic growth, a temporal pattern predicted to minimize generation time. Consistently, genetic manipulation of this developmental program reduced adult gonad size and delayed reproduction. Together, these findings reveal a temporally dynamic, rather than a static, resource allocation strategy that buffers reproductive timing against environmental stress through non-linear scaling of reproductive development.

## Results

### Reproductive timing scales with adult gonad size

To measure reproductive growth, somatic development, and reproductive timing, we used a *C. elegans* strain expressing GFP in the germline (*glh-1::T2A::sfGFP2*)^28^ and ubiquitously expressing a ribosomal protein functionally and endogenously tagged with mCherry (*rpl-34::RPL-34::mCherry*) ^29^. We acquired time-lapse movies of hundreds of individuals of this strain at 20-min resolution by automated live microscopy in agarose-based growth chambers^30^ (Fig. 1a). Measurement repeatability was high, with net experimental noise of 1% for total body size and 5% for the gonad size per individual and time point (Fig. S1).

**Figure 1.**
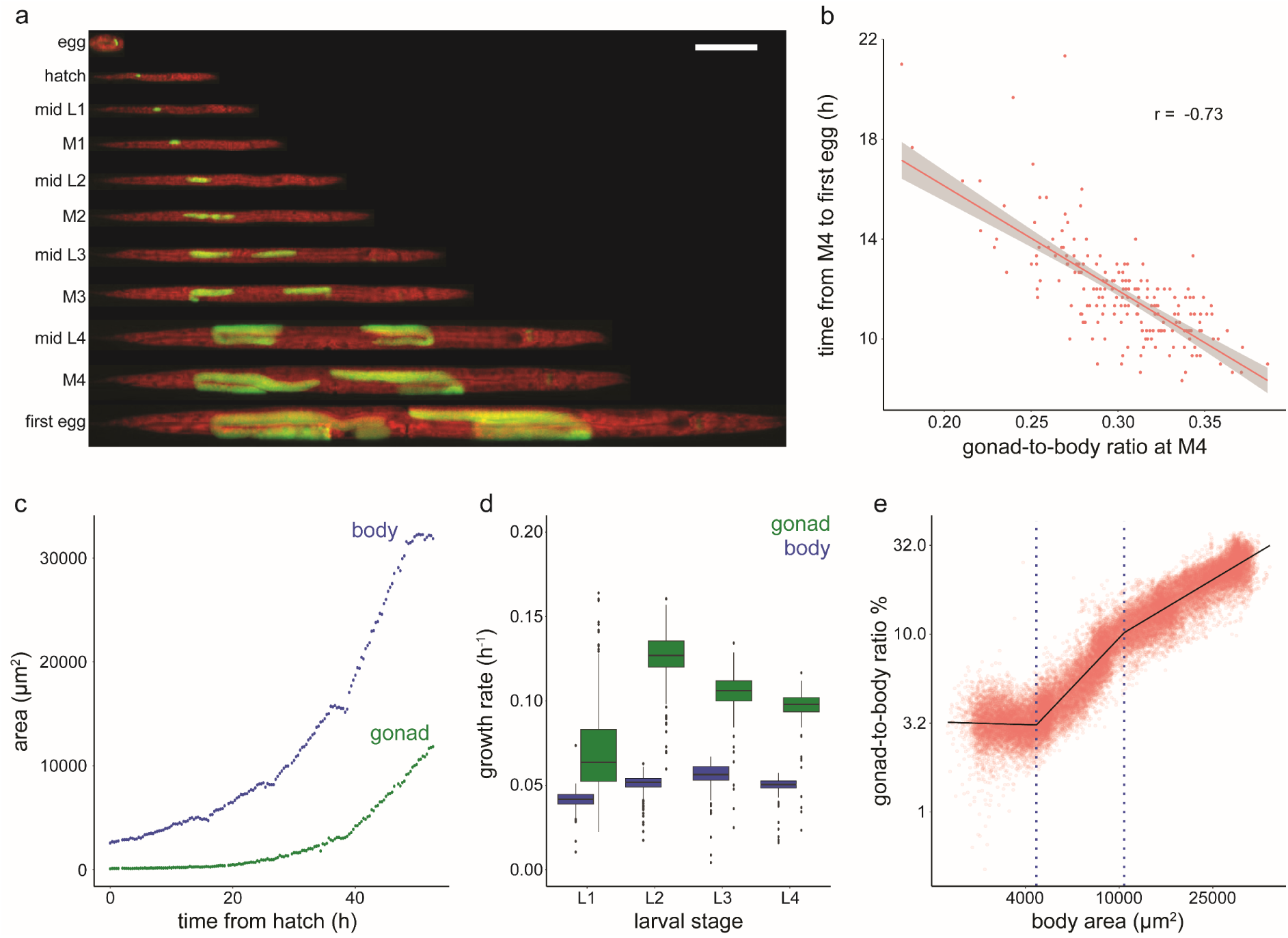
Gonadal growth is multiphasic. **a)** An individual animal imaged in agarose chambers at the indicated developmental milestones. Merged image of gonad in green (*glh-1::T2A::sfGFP2*) and total body in red (RPL-34:mCherry) is shown. Scale bar: 100 µm. **b)** Correlation of gonad-to-body proportion at M4 to time between M4 and onset of reproduction (r = −0.73 Pearson correlation coefficient, p < 10^-16^). **c)** Size trajectory of body and gonad from hatch to M4 for a single individual. Growth plateaus correspond to larval molts **d)** Growth rate (Δlog(area) / Δt) of body and gonads per larval stage. central line: median, box: interquartile range (IQR), whisker: ranges except extreme outliers (>1.5*IQR), circles: extreme outliers. **e)** Gonad-to-body proportions vs. body size in log-log scale. Each dot corresponds to one timepoint for a single individual. n of individuals = 259, n of measurements plotted = 39865, black line: fit of three-part segmented linear regression. dotted lines: separation of individual segments of the regression model.

Growth trajectories derived from 2D measurements were strongly correlated with volumetric measurements obtained by confocal microscopy (r = 0.992 and r = 0.984 for body and gonads, respectively, Fig. S2a-c), consistent with the near rotational symmetry of *C. elegans* ^30–32^. Unless otherwise noted, size measurements will refer to area in 2D, whereas validating volumetric measurements are mentioned explicitly. Gonad size will refer to the sum of both gonad arms.

On average, the gonads accounted for 3% (CI = ± 0.2%) of the total body size at hatching and expanded to 30.3% (CI = ±0.4%, CV = 10%) at the transition to adulthood, i.e. at the fourth larval molt (M4). Inter-individual differences in gonad-to-body proportions (gonad area normalized to body area) at M4 were negatively correlated with the time between M4 and the appearance of the first egg in the uterus (r = −0.73, Fig.1b). This association indicates a tight connection between gonad scaling at somatic maturation and subsequent reproductive timing.

### Gonadal growth sharply accelerates during development

To determine the relative dynamics of gonad and total body growth, we examined individual growth trajectories from hatch to M4. The body size increased near exponentially with time, except for transient plateaus during lethargus before each molt, as previously reported ^30^. However, whereas the molt-to-molt growth rate of the body (Δlog(A_B_)/Δt) was near-constant during development (Fig. 1d), the gonadal growth rate (Δlog(A_G_)/Δt) sharply increased between the L1 and the L2 stage (Fig. 1d).

Double logarithmic plots of the body size versus the gonad-to-body proportions revealed a sharp transition in the gonadal growth rate at a body size of twice the size at hatch (2.02-fold; 95% CI= ± 0.05). Below this size, the total body and the gonadal growth rate were nearly equal, and the gonad maintained an almost constant size proportion with mean proportions of 3.31% (CI = ±0.02%, slope = Δlog(A_G_/A_B_) / Δlog(A_B_) = 0.06 ± 0.004). Following growth acceleration above this threshold, gonad-to-body proportions increased continuously to 30.3% (CI = ±0.4%) by M4. Fits of a three-part segmented regression model to this data robustly identified the initial acceleration (slope: 1.67 ± 0.009), as well as a slight but reproducible slowdown in the gonad expansion during the L3 stage (slope: 0.95 ± 0.007). Both of these transitions were apparent in population-averaged and in individual-based trajectories (Fig. 1e; Fig. S3).

Volumetric measurements by confocal microscopy at higher resolution confirmed these results, showing that multi-phasic gonadal growth is not an artifact of spatial resolution or of 2D-projection (Fig. S2d). Although, on average, gonad acceleration occurred close to the first molt (M1), a significant spread for individual animals of up to 15h difference (Fig. S4) suggests that gonad acceleration is not directly related to cuticular shedding.

Together, we conclude that gonadal growth undergoes a transition from isometric to positive allometric growth shortly before the end of the first larval stage.

### Delay and acceleration of growth maintain adult gonad proportions under stress

Given the environmental plasticity of reproductive maturation in many species ^33,34^, we asked how the gonadal growth dynamics were affected by conditions that reduce the overall growth rate. Specifically, we investigated the response to dietary restriction (using the feeding-impaired *eat-2(ad1113)* mutant)^35^, and to osmotic stress (250 mM NaCl).

Although both conditions strongly reduced the total body growth rate and the adult body size (Fig. 2a, b), the proportion of gonad-to-body size at M4 remained nearly unchanged at 30.3% (Fig. 2c-e). Consistently, the time between the onset of egg production and M4 was not significantly affected by the stresses (Fig. 2f) and, as for unperturbed conditions (Fig. 1b), was strongly correlated with individual variation in gonadal size scaling (Fig. 2g). Thus, the endpoint of *C. elegans* gonad expansion and the correlated time between the last molt and reproductive onset are robust to changes in growth rates under stress.

**Figure 2.**
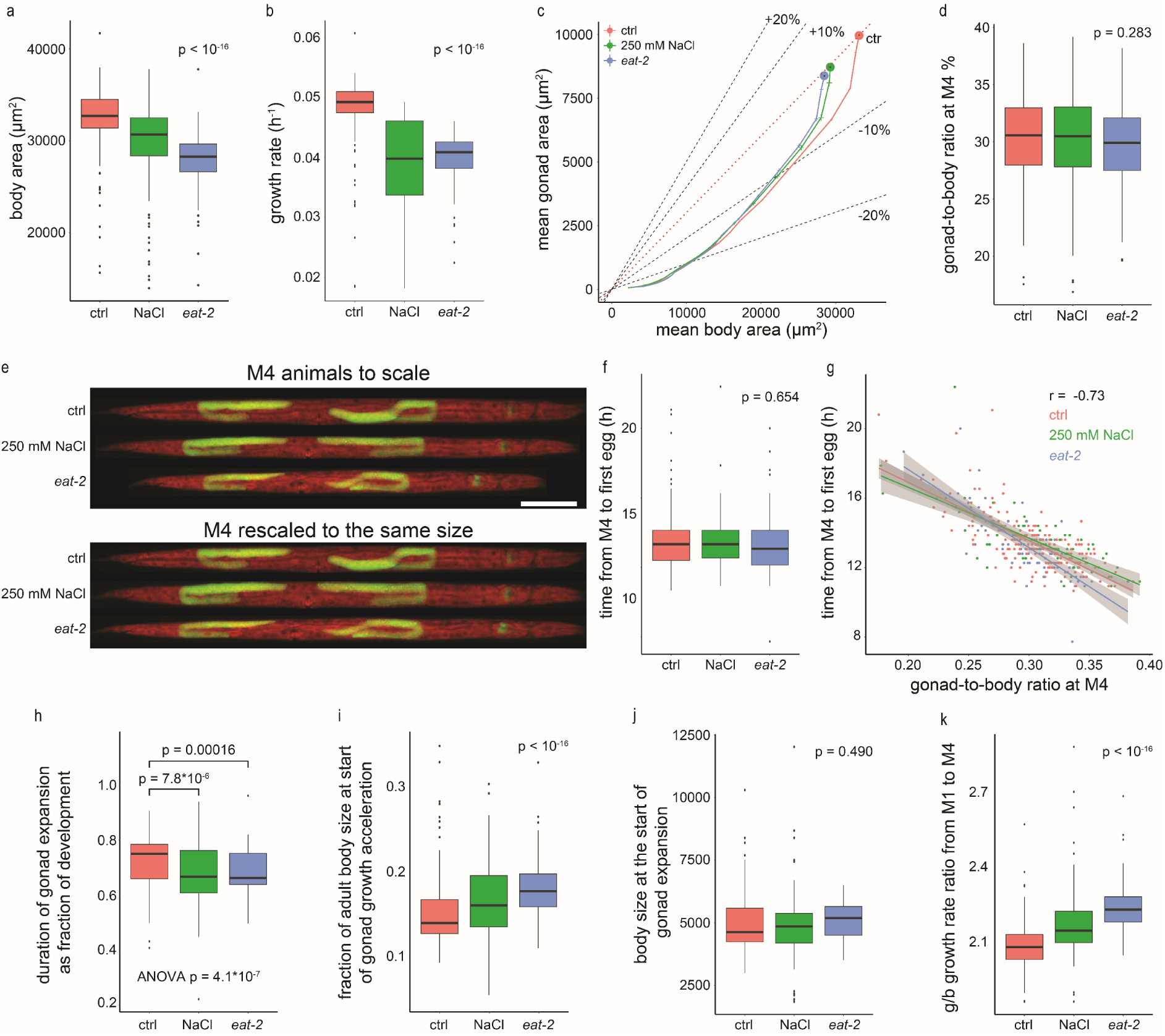
Environmental stresses modulate the gonad-to-body growth ratio to maintain adult gonad-to-body size proportions and reproductive timing. **a)** Mean body area at M4 for wild type with no salt added (ctrl, red), wild type with 250 mM NaCl added (NaCl, green), *eat-2(ad1113)* mutant with no salt added (*eat-2*, blue). **b)** Growth rate for the body from hatch to M4. **c)** Mean gonad size relative to mean body size with 95% CI. For each condition, longitudinal growth measurements starting at hatch and ending at M4 are plotted. **d)** Gonad-to-body ratio at M4. **e)** Images of computationally straightened worms from different conditions at M4 to scale and rescaled to the same body length. Scale bar: 100 µm. **f)** Time from M4 to first egg production. **g)** Correlation of gonad-to-body proportions with the onset of reproduction across individuals. h) duration of gonad expansion relative to duration of development. **i)** Fraction of the adult body size at the onset of the gonad expansion. **j)** Body size at the start of the gonad expansion. **k)** Ratio of gonad and body growth rates between M1 and M4. All panels except for f) and g): n =259, 96, 85 for control, 250 mM NaCl, and *eat-2,* respectively. For f) and g) n = 195, 73, 75 for control, NaCl, and *eat-2*, respectively. p-values are from ANOVA, except pairwise comparisons in h), which stem from Wilcoxon test. Box plots: central line: median, box: interquartile range (IQR), whisker: ranges except extreme outliers (>1.5*IQR), extreme outliers: circles.

However, unlike the final gonad-to-body proportions, the dynamics with which this endpoint was reached were highly plastic. Under both stresses, the transition to gonad expansion was delayed disproportionately relative to overall developmental duration, resulting in a compressed fraction of development devoted to gonad expansion (Fig. 2h, i). Notably, the threshold body size at which gonadal expansion started was unchanged by either of the two stresses (Fig. 2j), while the final adult body size was decreased (Fig. 2b). Consistently, the speed of gonad expansion relative to the total body was significantly increased under these conditions (Fig. 2k), as is necessary to maintain robust adult gonad-to-body proportions when the total body size is reduced (Fig. S5).

The timing of the first molt (M1), which occurs at a similar time as gonad expansion in unperturbed conditions, did not show the same behavior. Instead, the timing of M1 scaled proportionally with the developmental time in the *eat-2* mutant, and was shifted less strongly than gonad expansion under osmotic stress (Fig. S6). This temporal uncoupling between gonadal growth and molting further suggests that gonad expansion is not directly controlled by the molting timer or by the heterochronic pathway ^36,37^. Consistently, gonadal growth remained restrained during the first larval stage in *lin-14(n179ts)* mutants (Fig. S7), which precociously adopt L2 somatic cell fates during the first larval stage^38,39^.

Together, these data show that robust adult gonad-to-body proportions under stress emerge through plastic growth dynamics that temporally compress gonad expansion relative to the total duration of development.

### Acute phototoxic stress upregulates gonadal ribosomes levels and accelerates gonadal growth relative to the body

To ask how gonad scaling responds to acute, rather than chronic stress, we irradiated animals at three different doses (80s, 100s, and 120s) with near-UV light (405 nm) as L2 or L3 larvae, i.e., 20 hours or 30 hours after loading eggs into growth chambers.

Across all doses and timings, acute phototoxic stress slowed down body growth (Fig. 3a, b) and reduced the body size at M4 (Fig. 3c, d). As for continuous stresses, the gonadal growth rate increased relative to somatic growth after irradiation (Fig. 3e-h). As a result, while acute phototoxic stress during L2 or L3 impaired the overall growth rate and the size at M4 (Fig. 3c, d, Fig. S8), the gonad-to-body proportions at M4 remained close to unperturbed (Fig. 3g-j).

**Figure 3.**
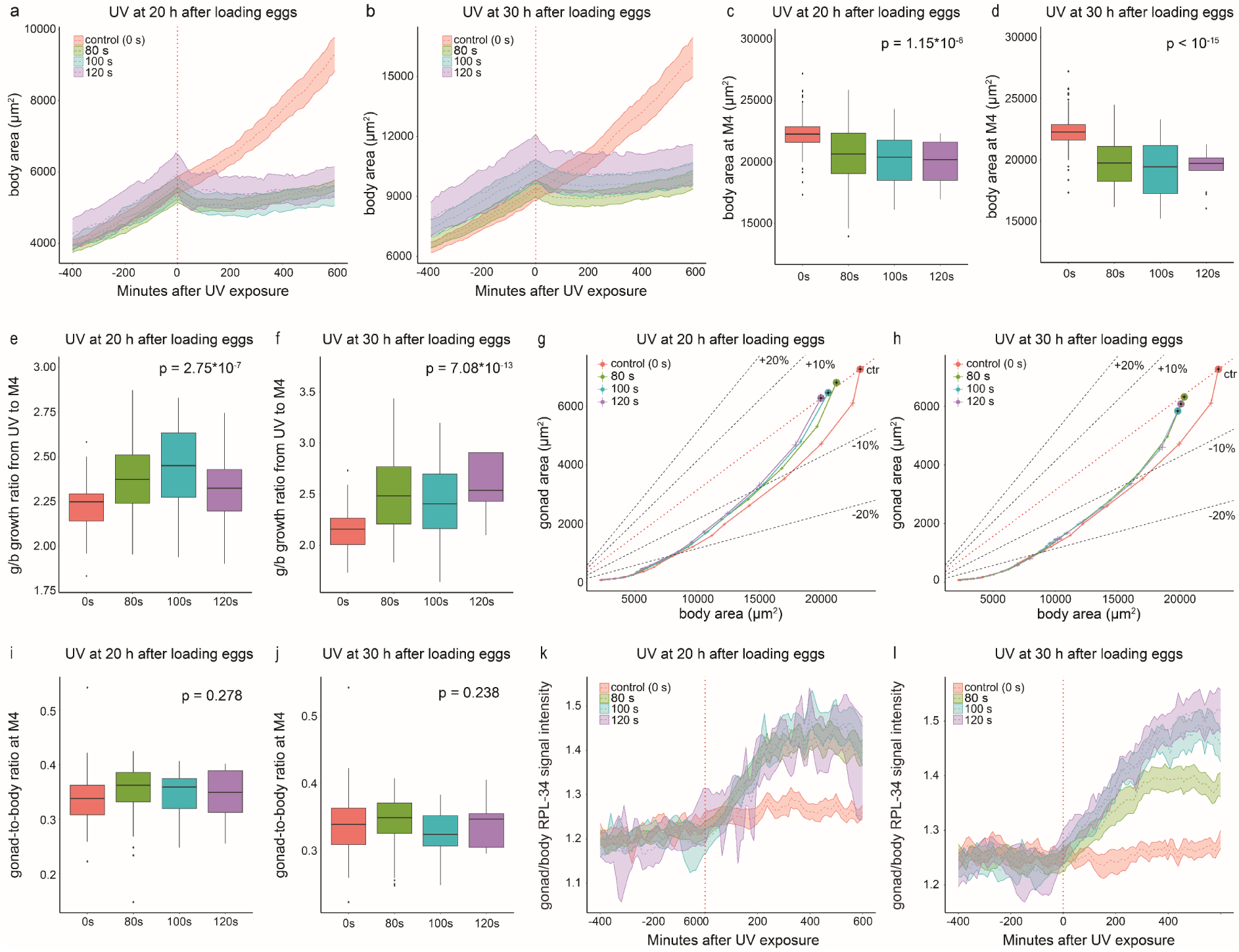
Gonadal growth accelerates in response to acute phototoxic stress to maintain adult gonad-to-body proportions. **a, b)** Body area before and after irradiation with near-UV light (405 nm) 20 hours (a) and 30 (b) hours after loading of eggs in chambers **c, d)** Body area at M4 after irradiation at 20 hours (c) and 30 hours (d). **e-f)** Ratio of average gonadal and somatic growth rates from time of irradiation to M4. **g, h)** Mean gonad size relative to mean body size with 95% CI. Dotted lines indicate gonad and body size with gonad-to-body proportions matching proportions observed in control conditions at M4 and indicated % deviation therefrom. **i, j)** Gonad-to-body ratios at M4. **k, l)** Mean signal of ribosomal subunit RPL-34:mCherry in gonads compared to signal in soma (body excluding gonads) before and after irradiation. For all panels with irradiation at 20h, n = 81, 76, 41, 14 for 0s, 80s, 100s, and 120s, respectively. Panels with irradiation at 30h, n =81, 76, 41, 14 for 0s, 80s, 100s, and 120s, respectively. The p-values: ANOVA.

Using a ribosomal protein RPL-34 functionally and endogenously tagged with mCherry ^29^, we next monitored ribosomal protein expression. Exposure to phototoxic stress was followed by an increase in the expression of the RPL-34:mCherry in the gonads compared to the surrounding somatic tissues (Fig. 3k-l), suggesting a tissue-specific regulation of translation capacity.

### A resource allocation model suggests a fitness benefit of biphasic delay-acceleration growth strategy

To ask if the observed gonadal growth dynamics can be explained by models of optimal life history plasticity, we analyzed a mathematical resource allocation model building on classic frameworks^5,6,40,41^ (Fig. 4a). In the model, total ingested nutrients (*N*) available for growth increase proportionally with the size (*s*) of the soma (*N* = *β* · *s*). Gonad and soma growth equally depend on nutrients following Michaelis-Menten (MM) kinetics, but only the soma contributes to future nutrient acquisition. Consistent with a metabolic cost of gonadal growth implicit in these assumptions, germline ablation by *glp-1* mutation accelerated development relative to wild type (Fig. S9).

**Figure 4.**
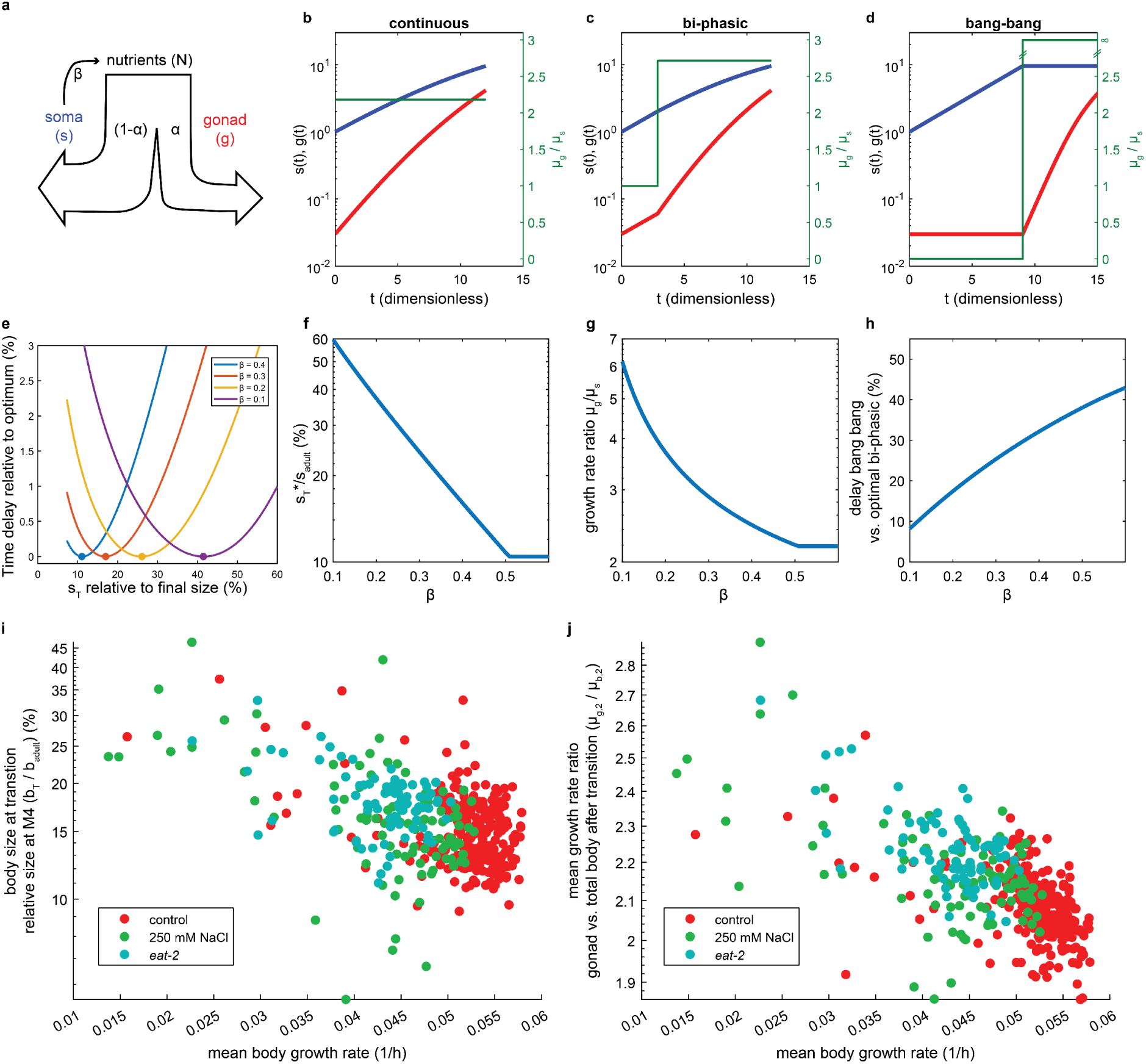
Resource allocation model suggests a fitness benefit of delay-acceleration growth strategy. **a)** Schematic illustration of the resource allocation model. **b)** Example model simulation of growth using α(t) that maintains a constant ratio of exponential growth rates of soma and gonad that reaches gonad-to-body proportions of 30.3% by adulthood. (Parameters: s_0_ = 1, g_0_ = 0.03, s_adult_ = 9.48, g_adult_ = 4.26, *β* = 0.33, μ = 0.33). Time refers to the nondimensionalized model, where t_dimensionless_ = t*μ. **c)** As b) but with α(t), such that *μ_s_* = *μ_gg_* for s < s_T_ = 2.0125 **d)** As b), but for α(t) = 0 when s < s_adult_ and α(t) = 1 when s = s_adult_, i.e., fully sequential growth of soma and gonad **e)** Relative delay in total time to adulthood as a function of s_T_ relative to development duration for optimal s_T_* for indicated values *β*. **f, g)** Relative size at transition to 2^nd^ phase s_T_*/ s (f) and growth rate ratio in 2^nd^ phase 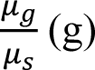 that minimize developmental time as a function of *β*. **h)** Relative delay of bang-bang control compared to bi-phasic growth with optimal s_T_*. **i, j)** Relation of experimentally measured ratio of body size at germline expansion onset to adult body size (i) and 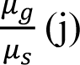, corresponding to model results shown in f) and g) versus the total body growth rate between M1 and M4 (Δlog(A)/Δt). Closed circles indicate individual animals, colors indicate conditions. Number of individuals n as in Fig. 2.

Nondimensionalization of time and nutrient uptake rate (Supplemental Text 1) yields a model with only one free parameter: the environment-dependent growth efficiency *β* (high *β* = high food uptake efficiency) (Eqs. 1-2). The relative nutrient allocation between soma and gonads is defined by a control function α(t) = N_g_/N_total 42_.

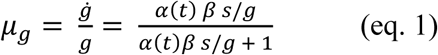

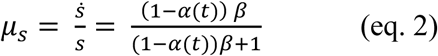

To approximate experimental observations (Fig. 1e), we used biphasic growth dynamics, where soma and gonads initially grow at equal rates (*μ_gg_*_,1_ = *μ_s_*_,1_). At a threshold body size s_T_, the gonads accelerate (*μ_gg_*_,2_ = *k* ∗ *μ_s_*_,2_), such that gonad-to-body proportions approach 30.3% once the adult body size is reached (Fig. 4b-d). For small values of *β*, a moderate delay in gonad expansion (Fig. 4c) shortened the time to reach the adult size compared to continuous exponential gonadal growth (Fig. 4b) or fully sequential soma and gonad growth (Fig. 4d, bang-bang control) (Fig. 4e-h). We next determined the optimal fraction of the final body size 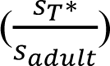 at which gonad acceleration onset minimizes the developmental time to adulthood. While 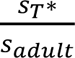 was independent of the adult body size (Fig. S10), 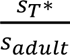 increased with lower nutrient uptake efficiency *β* (Fig. 4e, f). Thus, the model matches experimental observation (Fig. 2i) in predicting that under poor growth conditions it is preferable to delay gonadal growth acceleration. This consistency between model and experiment extended to the variation among individual animals, for which the growth rate was negatively correlated to the delay in the onset of gonadal growth acceleration (Fig. 4i, j), as predicted by the model (Fig. 4f, g).

We conclude that the experimentally observed plasticity of gonadal growth is consistent with a mathematical model predicting that time to maturity can be minimized by a relative temporal compression of gonad expansion under stress.

### Manipulation of the developmental clock reduces gonad-to-body size proportions and delays reproduction

To test whether uncoupling of gonadal from somatic growth impairs reproduction, we modulated the somatic developmental tempo by destabilizing the transcription factor BLMP-1 through auxin-induced degradation (AID). BLMP-1 is part of an oscillatory developmental clock that determines the duration of larval stages^43–45^.

As was previously reported^30^, destabilization of BLMP-1 sped up development relative to the growth rate, thereby reducing the final body size (Fig. 5a-b). However, unlike environmental stresses, BLMP-1 perturbation did not increase gonadal growth relative to the soma (Fig. 5c), such that gonad-to-body proportions at M4 were significantly reduced (Fig. 5d). Among individual animals, gonad proportions at M4 were negatively correlated to the delay in reproduction onset, following the same trend as unperturbed animals (r = −0.81). However, BLMP-1 AID animals were shifted towards smaller proportions and longer delays (Fig. 5e, f), suggesting that uncoupling of growth and developmental rates not only reduces the gonad size, but also negatively impacts reproductive timing. We note that the strain carrying the *blmp-1-aid* allele had slightly altered growth dynamics compared to untagged controls (Fig. 1), even in the absence of auxin. Importantly, however, since the experiments shown in Fig. 5f are internally controlled using the same tagged strain for treatment and control, we can exclude that the observed effect on the gonad size is caused by the *aid* tag alone.

**Figure 5.**
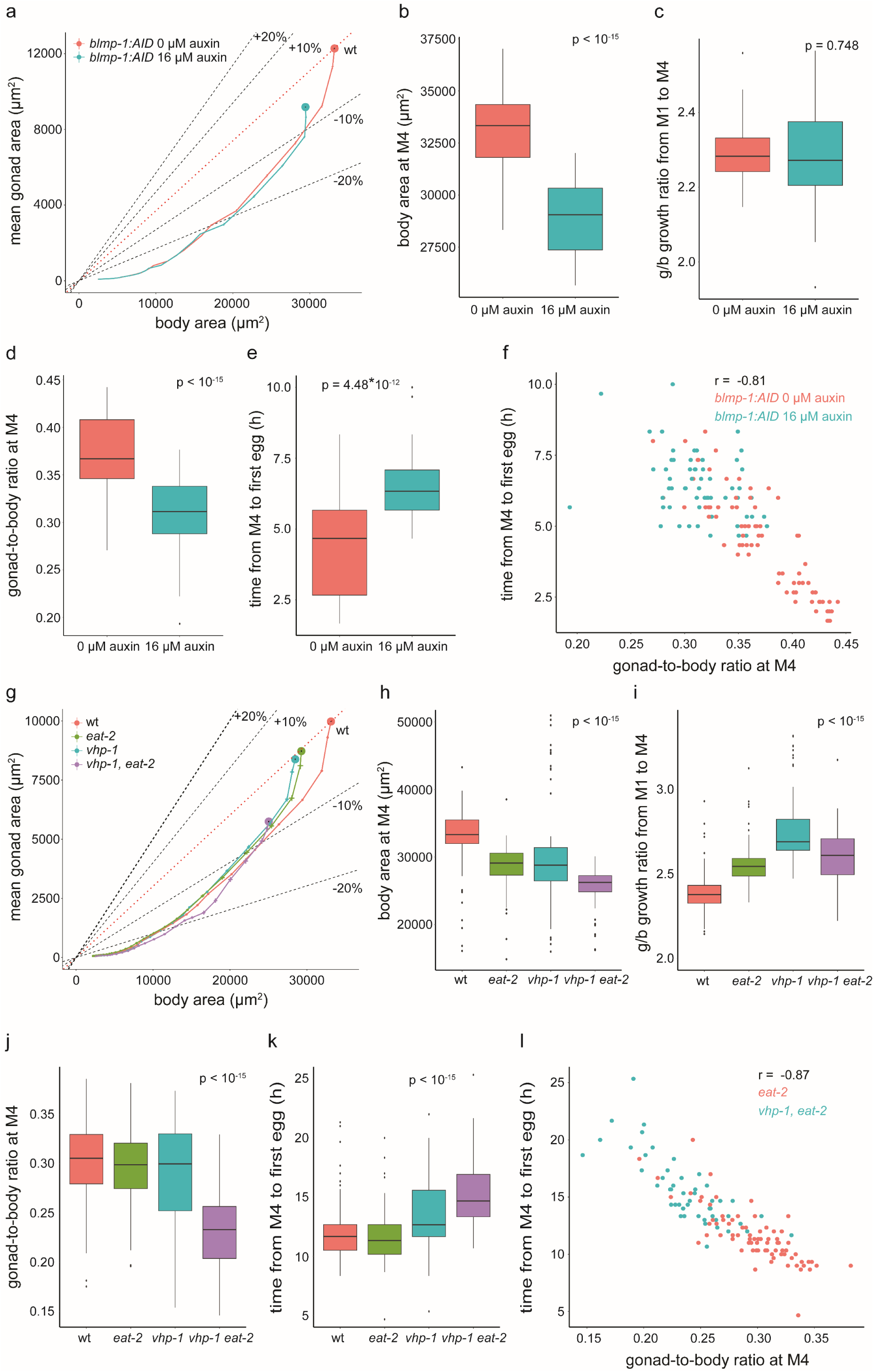
Altering gonad-to-body ratio at M4 delays reproductive onset. **a)** Mean gonad size relative to mean body size with 95% CI. **b)** body size at exit of M4 molt **c)** Ratio of gonad and body growth rates between M1 and M4. **d**) Gonad-to-body size ratio at M4. **e)** Time between M4 and first egg production. **f)** correlation of time between and M4 and first egg vs. gonad-to-body size ratio at M4 across individuals level for *blmp-1:AID 0 μM* auxin and *blmp-1:AID 16 μM* auxin. **g-l)** same as a-d, but for *eat-2*, *vhp-1(sa366),* and *vhp-1 eat-2* double mutant. **a-d)** n = 75, 61 for 0µM and 16µM respectively. **f-h)** n = 259, 85, 72, 64 for wt, *eat-2, vhp-1 and vhp-1 eat-2* double mutant respectively for i), n **=** 195, 75, 54, 50 for wt, *eat-2, vhp-1 and vhp-1 eat-2* double mutant respectivelly and for j), n = 75 and 50 for *eat-2* and *vhp-1 eat-2* double mutant respectively. The p-values: ANOVA.

We did not observe the same uncoupling between gonad and soma growth for a mutation of the non-oscillatory transcription factor *sma-4*, which similarly reduces body size (Fig. S11a) ^46^. Instead, *sma-4* mutants qualitatively matched the delay and acceleration of gonad growth observed in *eat-2* mutants and under environmental stresses (Fig. S11c, d) yet slightly increased the adult gonad-to-body proportions (Fig. S11b). Similarly, reproductive onset remained negatively correlated to gonad-to-body proportions (Fig. S11e) but was overall delayed compared to wild type (Fig. S11f), possibly due to pleiotropic effects of a strong body size reduction.

Together, these data show that, while reduced gonad-to-body proportions are not a general consequence of reduced body size, body size reduction by altering the developmental clock can uncouple somatic and gonad size and significantly affect reproductive timing.

### Stress-activated protein kinase (SAPK) signaling is required for the robustness of gonad scaling and reproductive timing under stress

To test whether stress signaling was involved in adjusting gonadal growth to environmental perturbations, we investigated a reduction-of-function mutant of *vhp-1(sa366)*, a dual-specificity phosphatase (DUSP) of the stress-activated protein kinases (SAPKs) p38 and JNK 47,48.

Similar to *eat-2* mutants and environmental stresses, the *vhp-1* mutation caused slow growth and reduced body size (Fig. 5g-h, Fig. S12) but had unchanged gonad scaling at M4 (Fig. 5j). This robustness suggests that *vhp-1* is not required for normal gonad proportions in the absence of stress. However, when *vhp-1* mutants were exposed to dietary restriction by *eat-2* mutation, gonadal growth was not sufficiently accelerated to attain correct proportions relative to the body size (Fig. 5i), resulting in undersized gonads at M4 (Fig. 5j). Consistently, the time from M4 to reproduction was prolonged under stress compared to wild-type animals (Fig. 5k). This effect was also observed for individual animals, where *vhp-1* mutants were shifted to smaller gonad proportions and slower reproduction (Fig. 5l), following an extension of the same trendline as observed for wild type animals. A similar, yet less pronounced, deviation in gonad-to-body proportions also occurred for *vhp-1* mutants exposed to osmotic stress (Fig. S13). We conclude that robust gonad scaling and reproductive timing under stress require intact *vhp-1*.

In summary, our data show that gonad scaling is robustly maintained across diverse environmental and genetic stresses but can be genetically uncoupled from somatic growth by destabilizing the oscillatory clock protein BLMP-1. The preservation of gonad-to-body proportions under stress is therefore not a passive outcome of reduced size or of slower growth. Instead, the inability of *vhp-1(sa366)* mutants to adjust gonad dynamics under stress supports an active, stress-induced regulatory mechanism involving p38/JNK signaling that coordinates the growth of somatic and reproductive tissues across different environments.

## Discussion

How reproductive timing is influenced by resource allocation between somatic and reproductive tissues is central to understanding mechanisms of fertility robustness and of life history evolution^1,5,6^. However, direct experimental quantification of resource allocation and its mechanistic control have lagged comparative and epidemiological studies of maturation timing. Organismal tests of predictions from life history theory, therefore, remain largely indirect. Our study addresses this gap by combining quantitative high-resolution measurements of gonad growth with an optimality framework and targeted genetic perturbations.

We show that *C. elegans* gonad growth is highly dynamic. Unlike somatic tissues, which grow continuously ^49^, the gonads accelerate their growth sharply near the end of the first larval stage (Fig. 1). Although environmental stress delays the onset of this acceleration relative to the total developmental duration, gonad-to-body proportions at adulthood remain preserved across environments. This robustness is due to the comparatively stronger acceleration of gonad growth following the delay (Figs. 2, 3), a pattern predicted to minimize maturation time by a resource allocation model (Fig. 4).

Our model suggests an explanation for how the observed growth dynamics could confer a reproductive advantage by minimizing the time to maturity. The model is based on two assumptions: first, although gonadal and somatic growth are metabolically equally costly, only somatic tissues contribute to nutrient acquisition. Second, physical constraints, such as enzyme saturation, impose an upper bound on the growth capacity of each tissue. Under these assumptions, delaying gonad expansion allows increased allocation of resources to somatic growth, thereby accelerating early development. The initially slow gonadal growth can later be compensated by increased nutrient allocation and faster growth. However, the maximal rate of this gonadal catch-up growth is constrained by enzyme saturation. When overall growth is slow, cells operate further below this capacity limit than under fast-growth conditions, allowing a greater relative acceleration of gonadal growth under stress. In the model, this faster acceleration enables a longer delay in gonad expansion without compromising final size proportions. These modelled relations may similarly account for the experimentally observed delay in gonad expansion under stress.

The preserved gonad-to-body proportions are strongly correlated with robust timing of first egg production under stress. Classic theoretical work shows that reproduction early in life contributes disproportionately to population growth ^50–52^, making the timing of reproductive onset a key determinant of fitness. Other traits, including reproductive rate, progeny quality, and brood size, are also likely to influence fitness under stress. The approaches developed here provide a framework to determine how these reproductive functions are coordinated and how potential trade-offs among them are balanced in future work.

The stress-induced relative temporal compression of gonadal growth observed here in *C. elegans* is reminiscent of epidemiological reports describing non-proportional sexual maturation dynamics in humans ^12,18,21–23^. Although such trends in humans are undoubtedly shaped by additional environmental and societal factors, our observations qualitatively parallel the relative expansion between early breast development (Tanner B2) and menarche under calorie-rich conditions ^21–23^, as well as reports of a relative compression of this interval under stress ^18^. It will be interesting to explore how these similarities relate to shared evolutionary pressures arising from constraints on resource allocation.

Together, our findings uncover previously unrecognized plasticity in the temporal control of gonadal growth. By dynamically reallocating growth resources between somatic and reproductive tissues, *C. elegans* ensure timely maturation across diverse environmental conditions. Given the parallels of the developmental dynamics between worms and humans, our study supports a central role of temporal growth allocation strategies in life history plasticity and advances the dissection of molecular mechanisms governing these reproductive growth decisions.

## Methods

### Strains used in this study

Animals were grown and maintained at 20°C on OP50-1 *E. coli* using nematode growth medium (NGM). The following strains were used in this study:

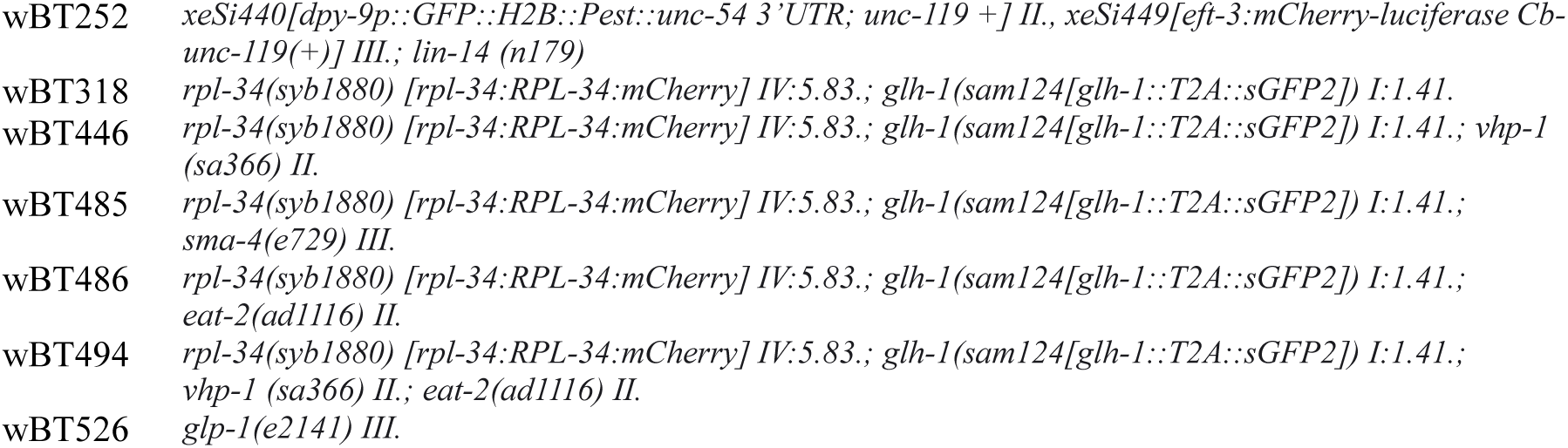

### Agarose chamber preparation and loading

Individual eggs were deposited in agarose-based chambers (4.5% agarose dissolved in S-basal), as previously described ^49^ and imaged every 20 minutes at 20°C. In brief, an inverse template made from polydimethylsiloxane (PDMS) was used to pattern an agarose gel with wells. Wells were filled with bacterial strain OP50-1. Individual eggs were manually placed into each well and chambers were sealed by inverting the agarose onto a microscopy-grade gas permeable polymer mounted on a 3.5cm dish (ibidi). Subsequently, the agarose-based array of chambers was surrounded by 3% low melting temperature agarose, dissolved in S-basal, and covered with PDMS to prevent drying of the sample. Chamber dimensions were 600 × 600 × 20 µm, except for the acute phototoxic stress experiments, where chambers with dimensions of 750 × 750 × 45 µm were used to reduce escaping of animals agitated by irradiation. For chronic stress experiments, imaging was performed on Squid microscopes from Cephla Inc. (https://cephla.com/), equipped with a Blackfly S CMOS camera (BFS-U3-63S4M-C, Teledyne Vision Solutions) and a 10x / N.A. 0.4 Olympus objective ^53^. Acute phototoxic stress experiments were performed on a Nikon Ti2 microscope equipped with a Hamamatsu ORCA Flash 4 camera and a Crest V3 spinning disk unit operated in widefield mode. A Celesta light engine was used for imaging and for phototoxicity stress illumination.

### Image analysis and calculation of volumes and growth rates

The outline of the body and gonads was determined by a UNET++ deep learning algorithm trained on a manually curated and adjusted segmentation mask created by the Segment Anything for Microscopy (μSAM) ^54^. Segmentations of body and gonads were subsequently straightened using the body outline as a template, as previously described ^30^. The full image analysis pipeline used in this project is accessible at the GitHub repository: https://github.com/spsalmon/towbintools_pipeline.

A random forest-based classifier^30^ was used to determine the time of hatching and to identify time points at which straightening failed (e.g., due to self-touching). Molts were detected from body volume trajectories as previously described ^30^.

### Stress induction

Chronic osmotic stress was applied by increasing the NaCl concentration in agarose chambers by 250 mM in both the high-melt and in the low-melt agarose fractions. Animals that did not reach M4 during the experimental time were not included in the analysis.

### Phototoxic stress

Each animal was exposed to 80s, 100s, or 120s of irradiation by the 405nm laser line of the Celesta light engine (Lumencor) at 100% power (∼500mW nominal fiber output). Exposure times were selected so that the highest dose caused 50% mortality. Only the growth trajectories of worms that reached M4 were included in the final analysis.

### Estimation of net instrument noise

To estimate net measurement noise in body and gonad size, time series of 10 images acquired with ∼0.3 frames per second were obtained for each of 10 animals using an independently executed separate autofocus for each acquisition. These measurements were repeated at 30-minute interval for each of the 10 individuals. The measurement error was determined as the median CV among repeated measurements at each time point (Fig S2), assuming that the true size does not change on the short time scale of 3 seconds.

### Comparison of area and volume measurements

To compare area measurement with volumetric measurements, two separate experiments were conducted to measure body and gonadal volume (Fig. S1).

First, we performed longitudinal experiment measurements of five individuals in agarose chambers as described above, except that z-stackes with 1µm spacing were acquired using a Ti2/Yokogawa W1 spinning disk confocal microscope and a 60x oil objective. Chamber size was adjusted to 230 × 230 × 10 µm to match the smaller field of view at this magnification. Chambers of this size allow completion of the second larval stage (Fig. S3a-c).

Second, mixed cultures with animals from L1 to L4 stages were grown on standard NGM plates, immobilised on 2% agarose pads using Levamisole (10mM), and imaged using a Ti2/Yokogawa W1 spinning disk confocal microscope and a 60x oil objective (Fig. S3d) as z-stacks with 1µm spacing. Individual fields of view were computationally stitched prior to measurement of gonad and body size.

### Auxin treatment

Auxin (IAA, Sigma) solutions were prepared on the day of the experiment as a 400x stock in EtOH and subsequently diluted to the indicated concentration in agarose to a final EtOH concentration of 0.25% immediately prior to use for micro chamber assembly.

### Measurement of reproductive timing

To measure reproductive timing, we have manually annotated images showing the first appearance of an egg inside the worms’ bodies (See Fig. 1a).

### Measurement of growth rates

The average growth rates, e.g., from M1 to M4, were calculated as the difference between the natural logarithm of the volume at the beginning and the end of the period divided by the larval stage duration 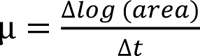.

### 3-segment linear regression

To quantify changes in gonad-to-body scaling during development, we performed segmented (piecewise linear) regression with up to two breakpoints (three linear segments) per individual. For the purposes of fitting *x* = *log*_10_ and All analyses 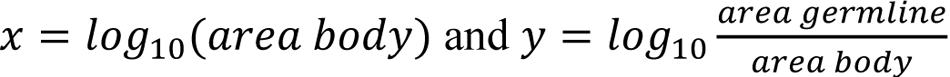 were conducted in R using the package segmented.

## End notes

### Author Contributions

**P.L.** formulated the research question, performed the experiments, analyzed data, and drafted the manuscript. **I.G.** conducted critical preliminary research using the *vhp-1* mutant, **S.P.** established image analysis workflows, **B.D.T.** supervised the project, guided experiments and data analysis, designed the model, and edited the manuscript. All authors reviewed the manuscript.

## Supplementary Information is available for this paper

### Code availability

The image analysis pipeline used in the study is accessible at the GitHub repository: https://github.com/spsalmon/towbintools_pipeline. The code for model simulations is provided as a supplemental data file.

