## Supplemental figures and text for "Temporal compression of gonadal growth preserves reproductive timing and soma-to-germline scaling under stress"

##### **Content:**

- Supplemental Figures 1-13
- Supplemental Text 1

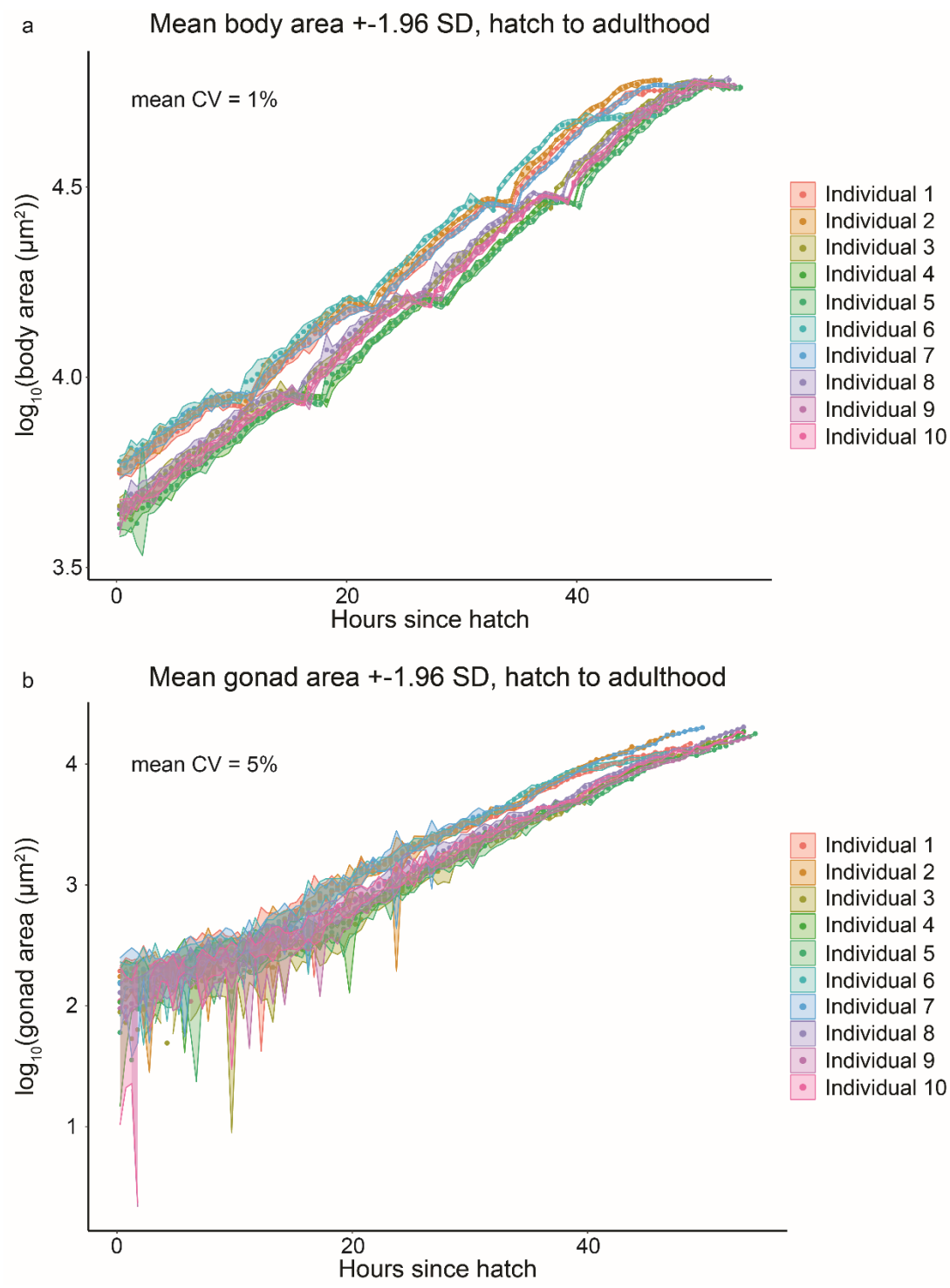

13 **Supplemental Figure 1. Raw measurement error (CV) of gonad and body area at individual time**
14 **points.** n = 10 individuals were imaged in agarose chambers at 20-min intervals, and 10 consecutive
15 images were acquired at each time point. A separate auto focus was performed for each acquisition.
16 Plots show measurements of the individuals distinguished by different colours. Circles indicate the
17 mean of 10 repeated measurements. The shaded area is $\pm 1.96 \times \text{SD}$ among the repeated measurements.
18 When the shaded area is invisible, the error is smaller than the marker

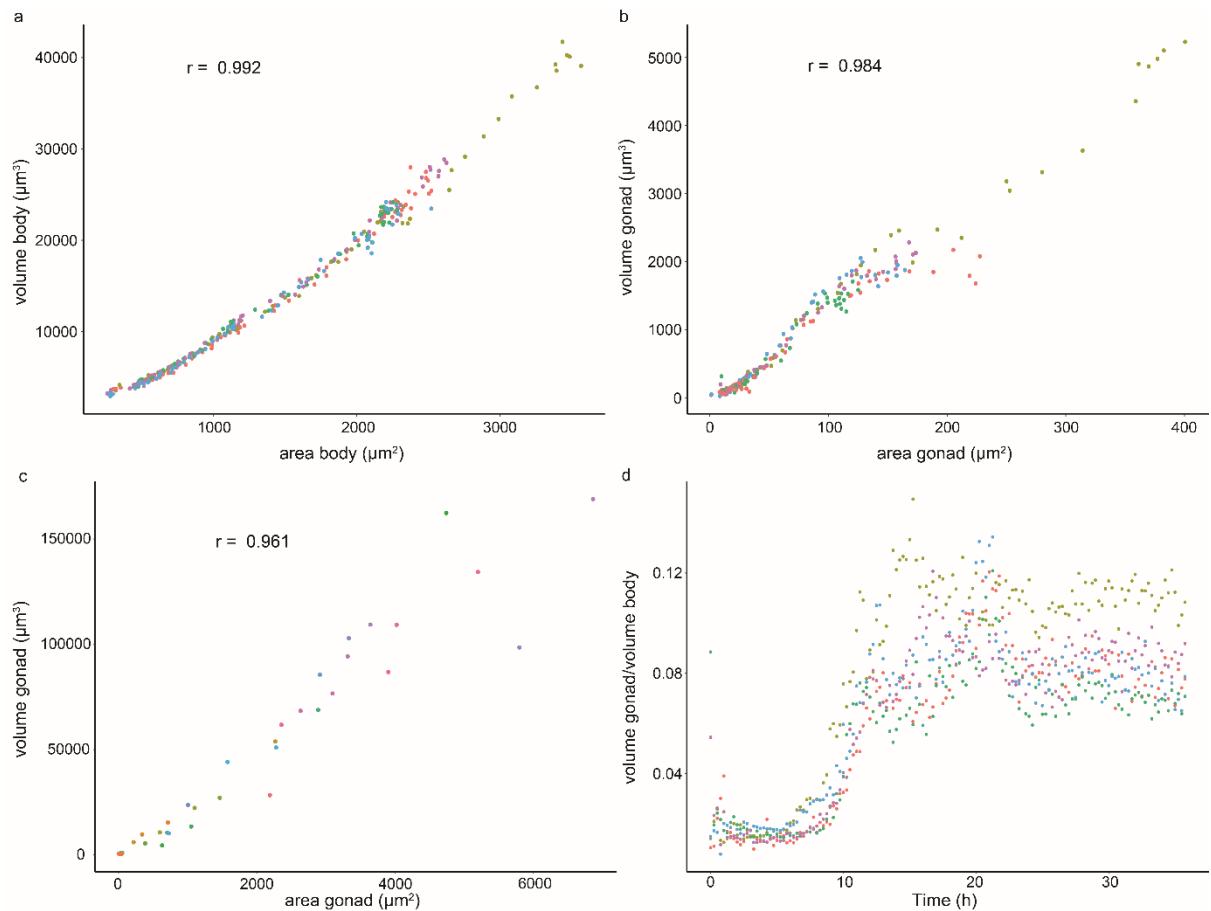

**Supplemental Figure 2. Validation of area-based measurement by volumetric confocal microscopy.** **a,b,d)** Longitudinal growth measurement in agarose chambers using 60x oil objective and a spinning disc confocal microscope **a)** Comparison of body volume determined from 3D confocal z-stacks with body area measurements from 2D maximum intensity projections of these same z-stacks. Each colour represents a different individual, $n = 5$. Small chamber size needed to accommodate the reduced field of view of a 60x magnification objective allowed imaging until M2, after which animals arrested due to food clearance. Only measurements from the growing phase are plotted. **b)** as a), but comparison between gonad volume and area. **c)** Comparison of volumes and areas of gonads in animals at different stages of development from L1 to M4 obtained by single snapshot imaging of immobilised animals on agarose pads using 60x oil objective, $n = 37$. **d)** Change in gonad-to-body proportion during the longitudinal experiment used in a) and b) computed from volumetric measurements. A rapid increase in gonad-to-body proportions around 10h is observed, consistent with 2-dimensional measurements shown in the main figures. Plateau after 20 hours corresponds to growth arrest due to food clearance in small chambers.

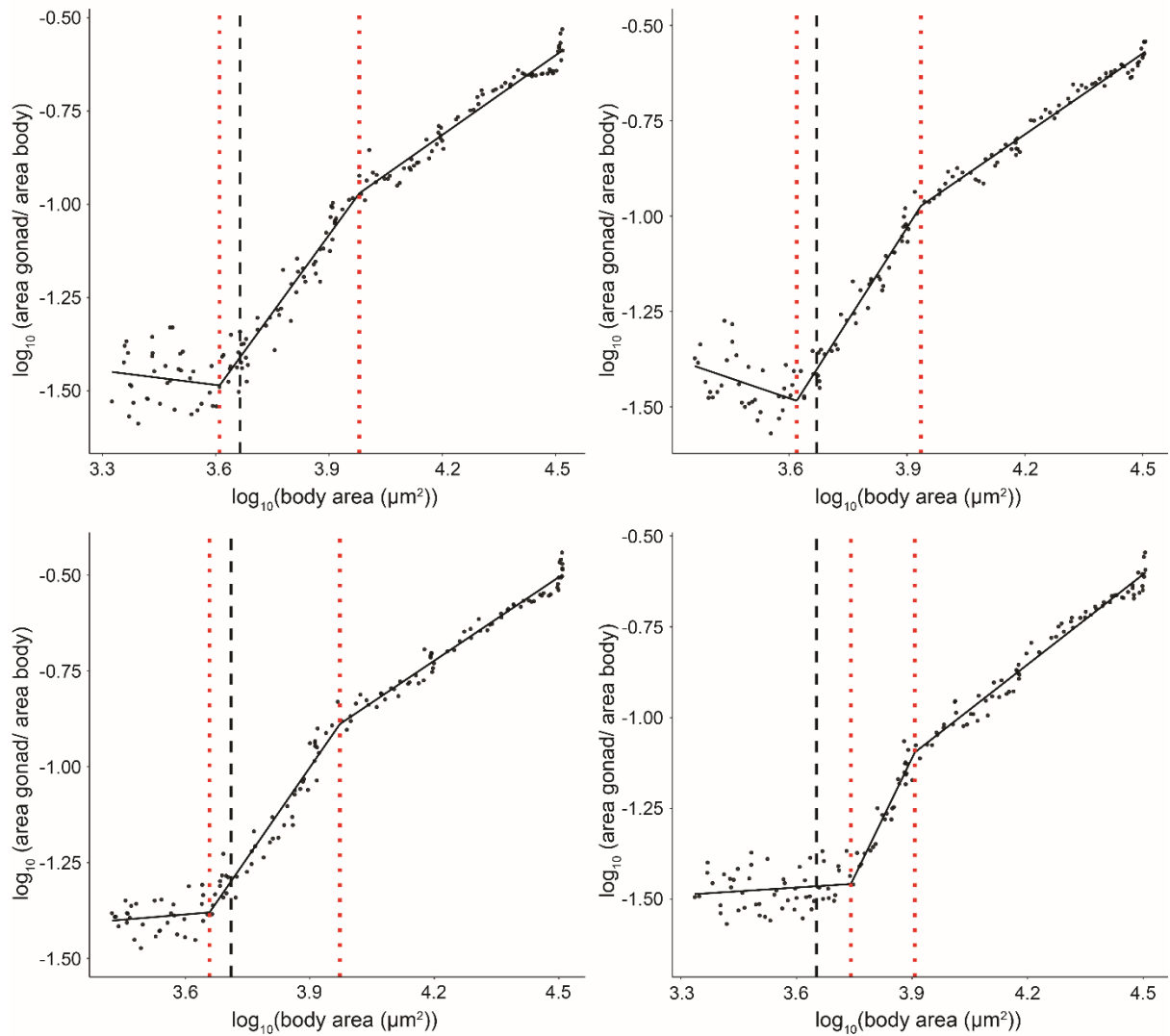

**Supplemental Figure 3. Examples of growth trajectories for individual animals with segmented regression.** Each dot represents a single measurement of an individual animal. The black line shows segment regression. Red dotted lines show boundaries between segments. The black dashed line shows the exit of the M1 molt.

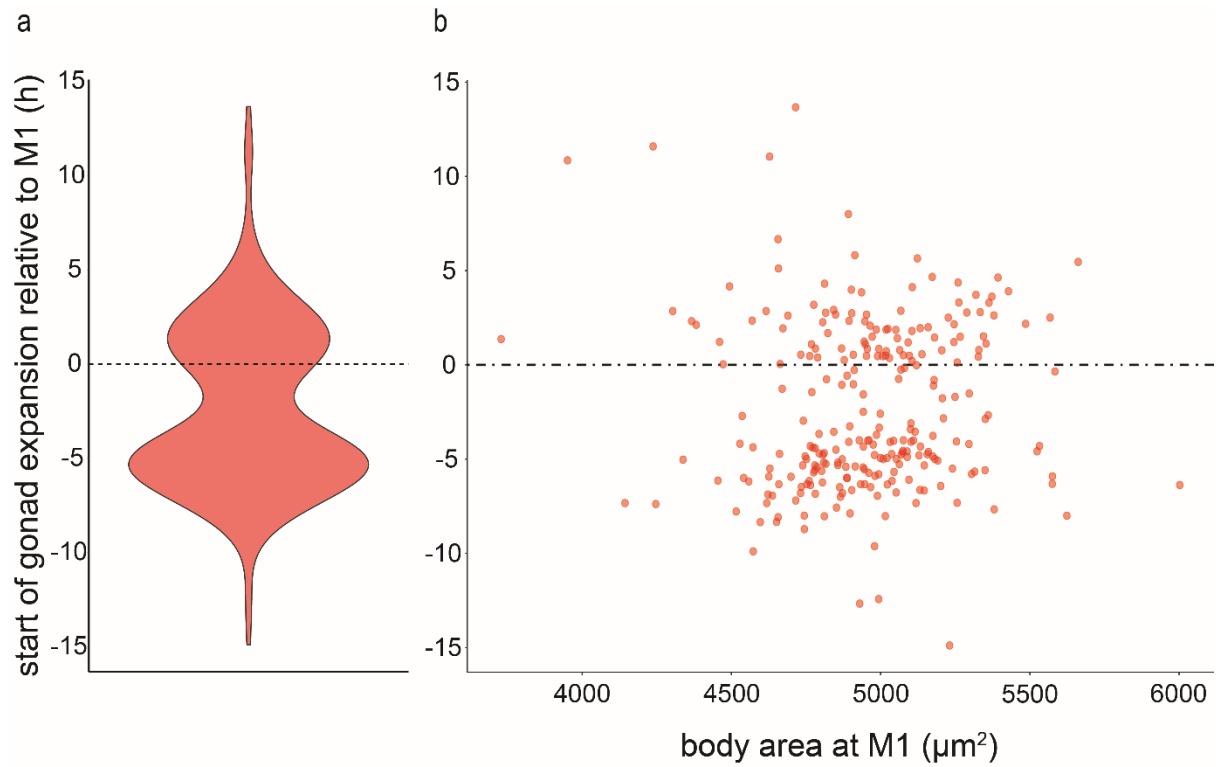

**Supplemental Figure 4. The onset of the gonad expansion does not coincide with M1 ecdysis. a)** Violin plot of the difference between the onset of gonad expansion and the ecdysis of M1 molt. **b)** Plot of individual worms by size at M1 and the difference between the onset of gonad expansion and the ecdysis of M1 molt. Sparsely populated region below 0 corresponds to lethargic phase prior to ecdysis. n = 259.

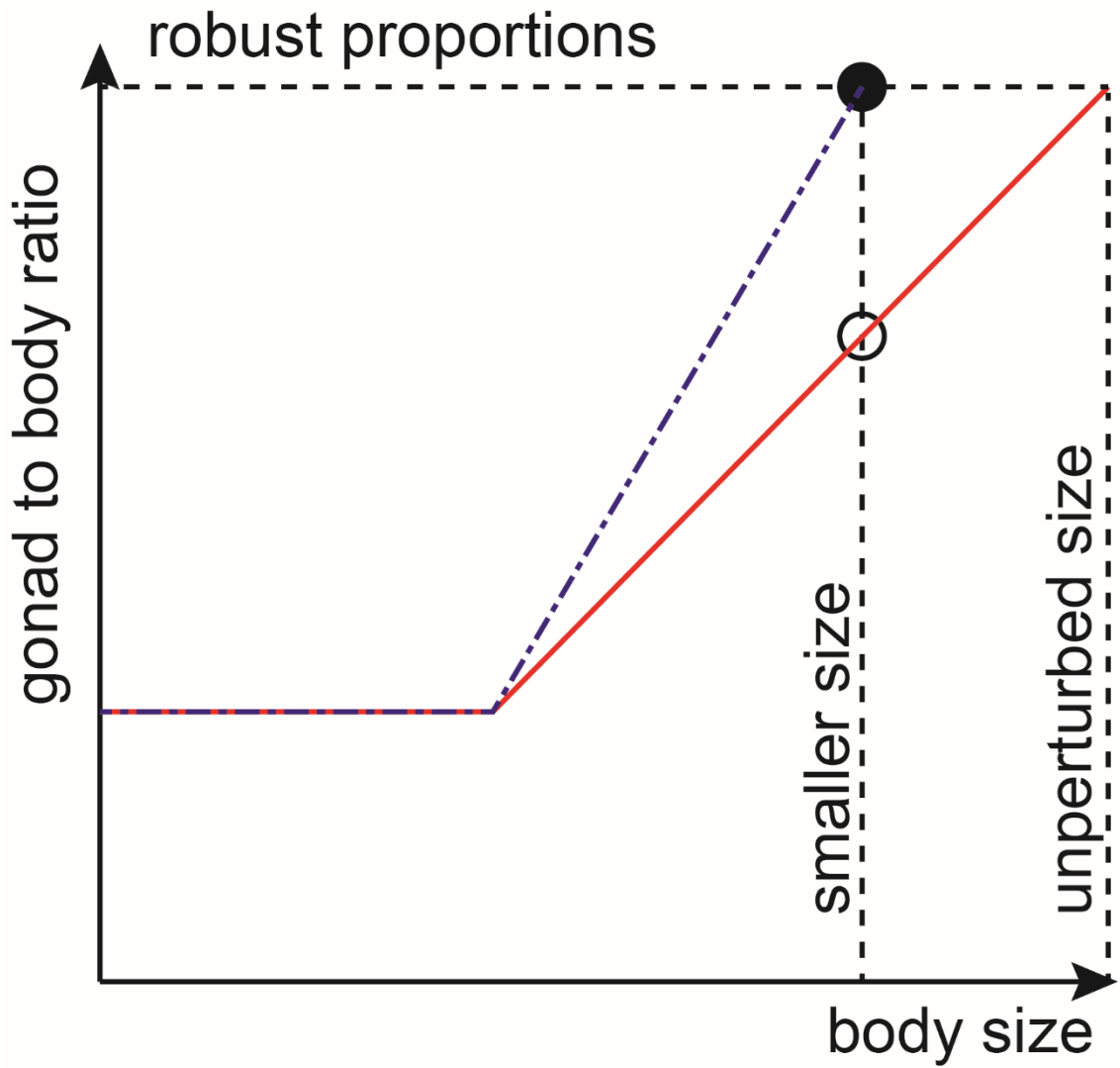

57

58 **Supplemental Figure 5. Schematic illustration of effect of adult body size on gonad-to-body**
59 **proportions.** If the adult body size is reduced and the size at which gonad expansion starts remains
60 unchanged, the rate of gonadal growth relative to somatic growth must be increased to keep the final
61 proportions constant. Open circle indicates gonad-to-body proportions when adult size is reduced
62 without changing the gonad-to-body growth rate ratio (red). Filled circle indicates robust proportions
63 when gonad growth is appropriately accelerated relative to the body (blue).

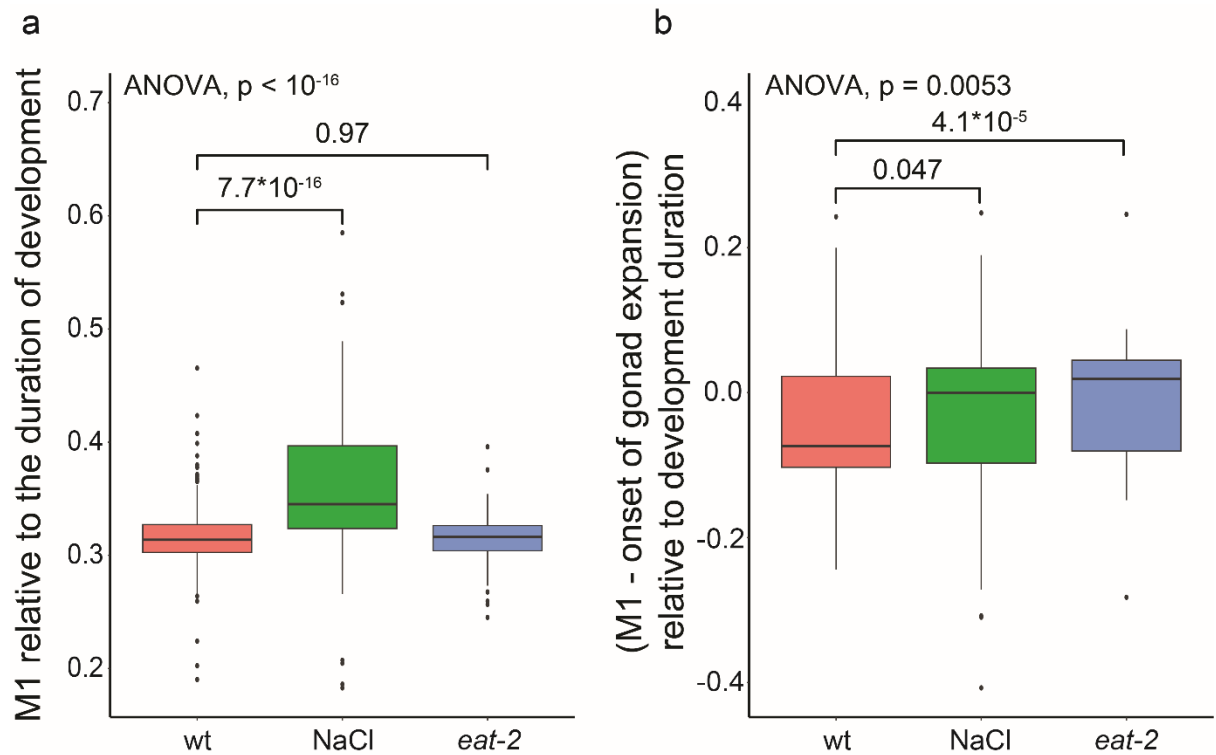

**Supplemental Figure 6. M1 ecdysis and the onset of gonad expansion scale differently with stress.**
**a)** Time of M1 onset as a fraction of the total duration of development. **b)** difference between M1 ecdysis and the onset of gonad expansion normalized to the total duration of development. P-value group comparisons: Two-sided (a) or one-sided (b) Wilcoxon rank test. $n = 259, 96, 85$ for control, 250 mM NaCl, and *eat-2*, respectively.

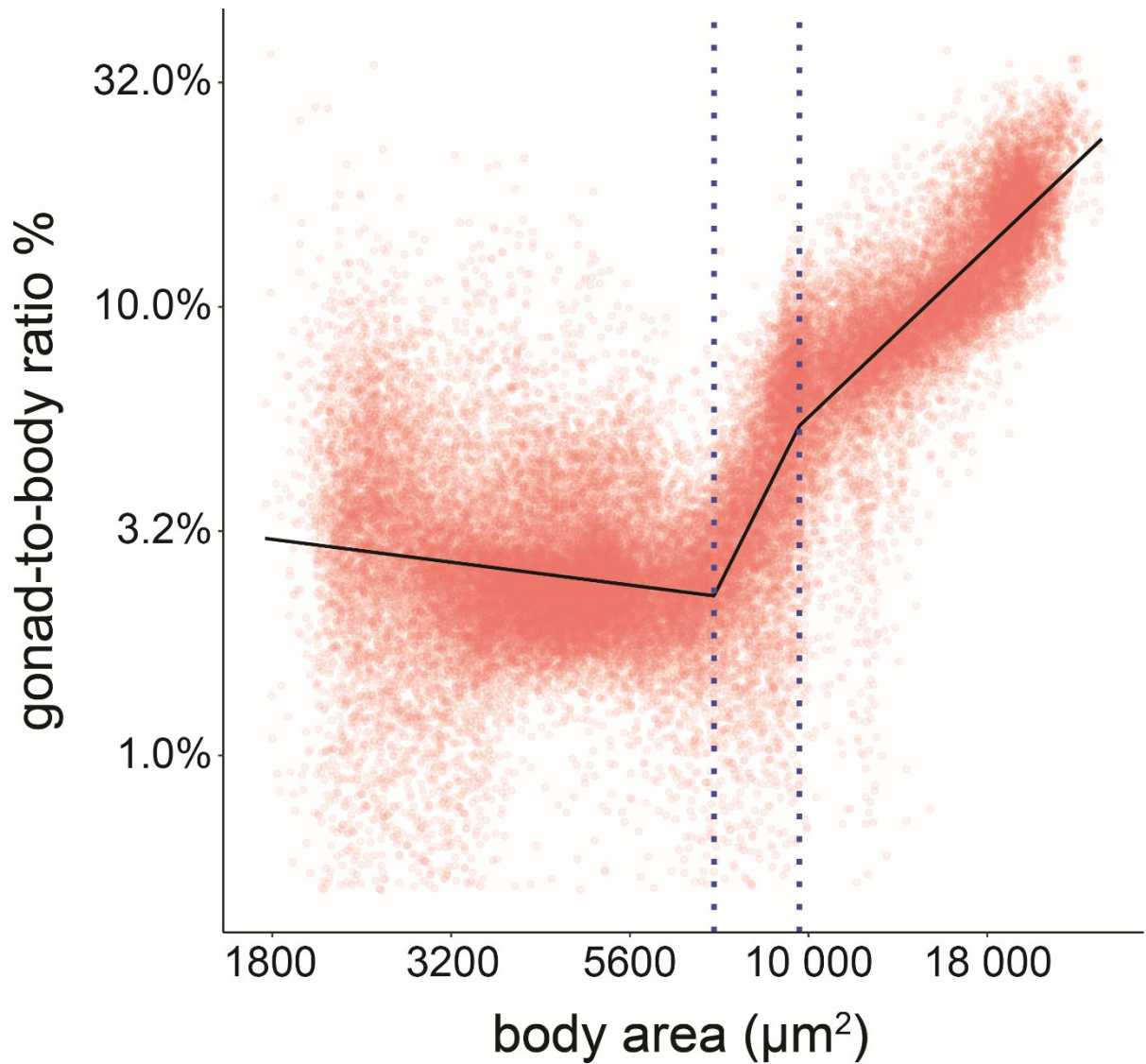

**Supplemental Figure 7. Gonadal growth remains restrained in a heterochronic mutant skipping** **L1 stage.** Gonad-to-body proportions vs. body size of *lin-14(n179ts)* reduction of function mutant grown at restrictive temperature (25°C) shown in double logarithmic scale. Each dot corresponds to one timepoint for a single individual. black line: fit of three-part segmented linear regression. n of individuals = 229, dotted lines: partition between individual segments of the regression model. The mutant shows constrained gonadal growth during early development, similar to wild type, despite adopting somatic L2 cell fates in the first larval stage after hatching. For this experiment, a strain expressing GFP in somatic tissues was used (*eft-3p:gfp*) and the gonad size was determined by a UNET++ model that recognizes the non fluorescent regions, i.e., the germ cells not expressing GFP.

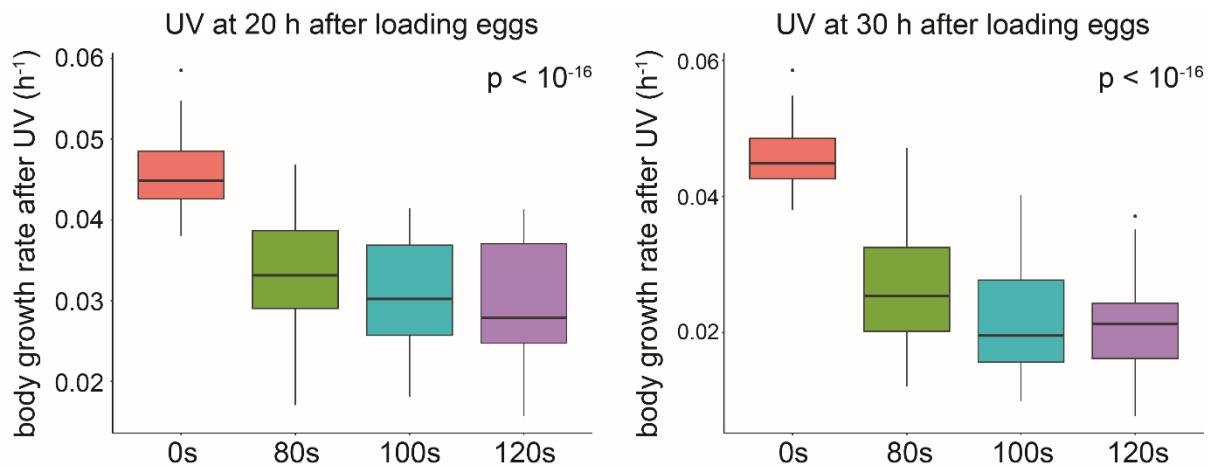

**Supplemental Figure 8. Body growth slows down in response to acute phototoxic stress.** Boxplots shows growth rate from UV irradiation to M4. For irradiation at 20h, $n = 81, 76, 41, 14$ for 0s, 80s, 100s, and 120s, respectively. For irradiation 30 h after the start of the experiment, $n = 81, 76, 41, 14$ for 0s, 80s, 100s, and 120s, respectively. The p-values: ANOVA.

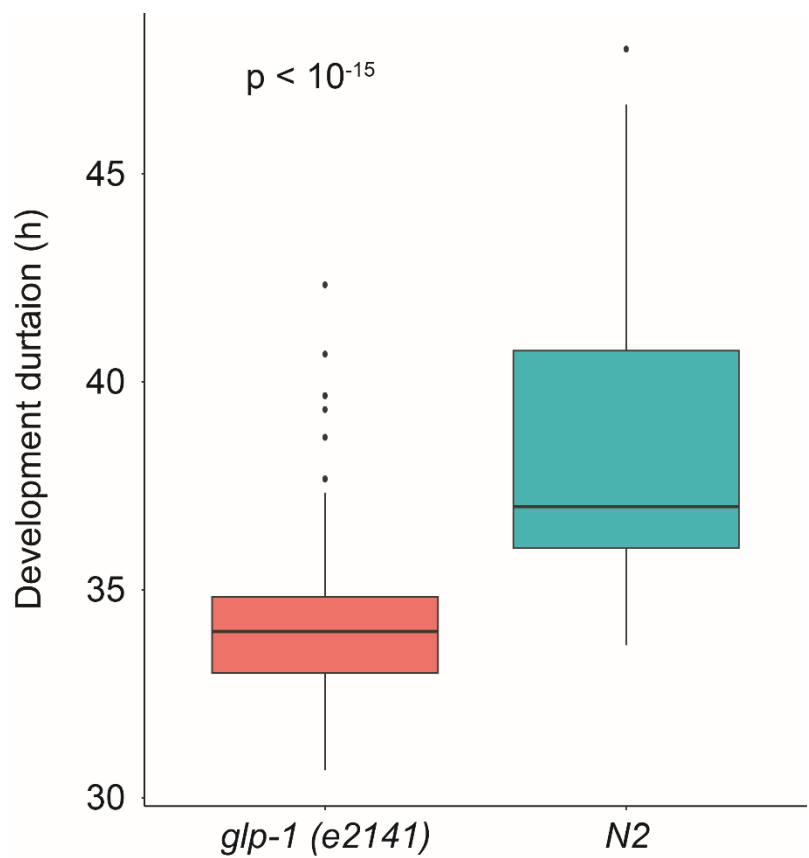

**Supplemental Figure 9. Genetic germline ablation by *glp-1* mutation shortens development.** Boxplot showing time from hatch to exit of M4 molt at 25°C, $n = 135$ and 144 for the temperature-sensitive *glp-1(e2141)* mutant and wild type (N2) respectively. The maternal generation of both strains was cultivated at 15°C. The p-values: ANOVA.

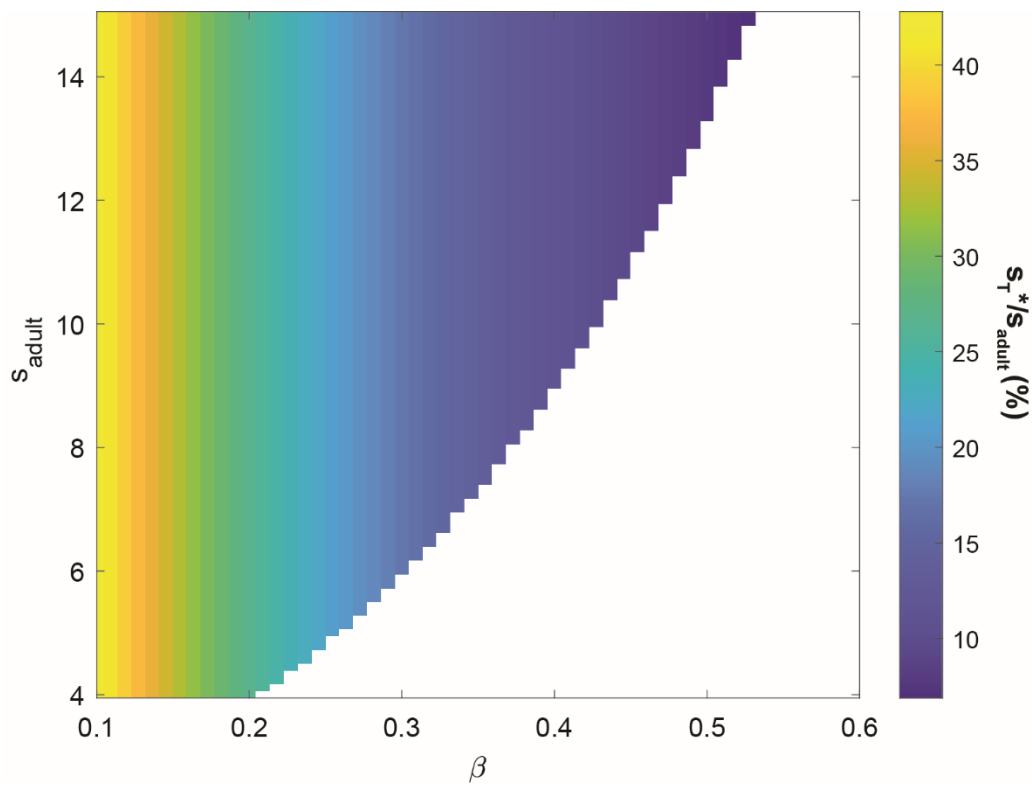

95

96 **Supplemental Figure 10. Dependence of optimal onset of gonad expansion on nutrient uptake**
97 **efficiency and adult body size.** Heatmap of the optimal fraction of adult size ($s_T^*/s_{adult}$) for gonad
98 expansion onset that minimizes time to adulthood as a function of nutrient uptake efficiency $\beta$ and the
99 final adult size, computed from model shown in main text. The white area indicates the parameter space
100 in which rapid gonad expansion from hatching (no delay) minimizes time to maturity.

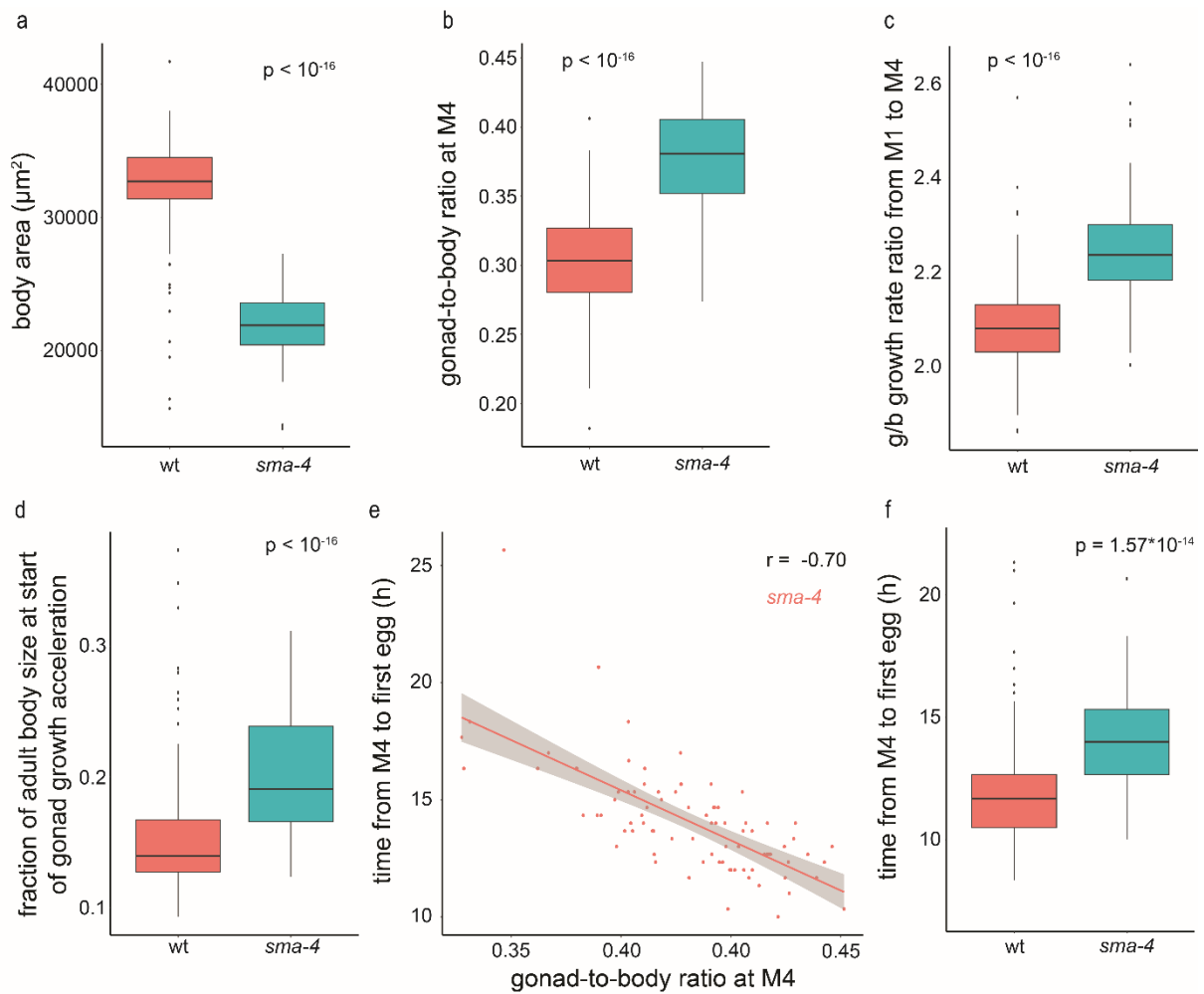

**Supplemental Figure 11. The *sma-4(e729)* mutation leads to a smaller size, accelerated gonadal growth, and increased gonad-to-body proportions.** **a)** Mean body area at M4. **b)** Gonad-to-body size ratio at M4. **c)** Ratio of gonad and body growth rate measured between M1 and M4. **d)** Fraction of the adult body size at the onset of the gonad expansion ($S_T/S_{\text{end}}$). **e)** Correlation of gonad-to-body proportions with the time between M4 and first egg production for individual animals. **f)** Time from M4 to first egg production. For a-d), $n = 259$, 90 for wt and *sma-4*, respectively. For e-f), $n = 195$, 84 for wt and *sma-4*, respectively. p-values: ANOVA.

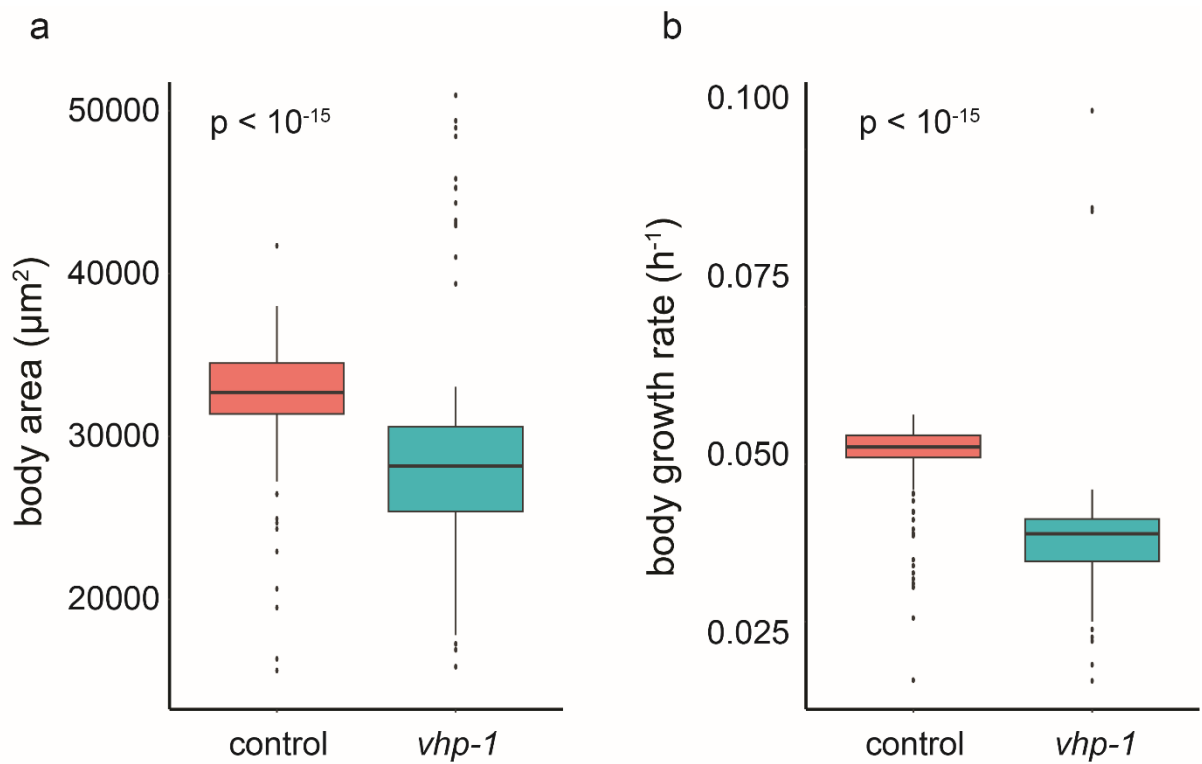

**Supplemental Figure 12. *vhp-1* (*sa366*) mutant has decreased adult size and slowed down growth.**

**a)** Body area at M4. **b)** Body growth rate measured from M1 to M4, n = 259, 72 for control and *vhp-1*.

The p-values: ANOVA.

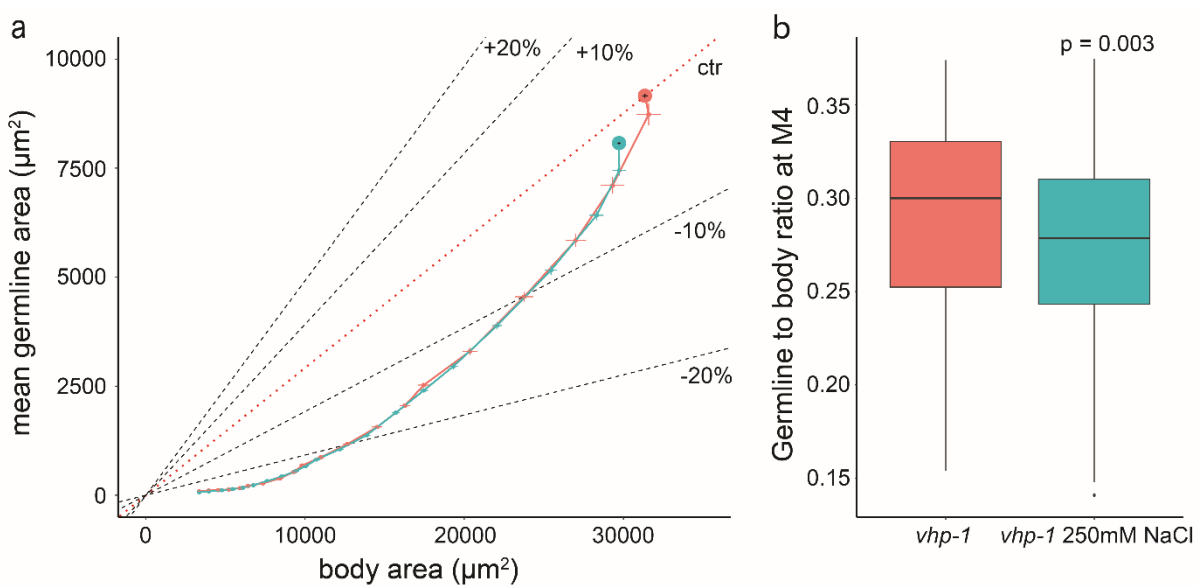

**Supplemental Figure 13. Gonad-to-body proportions of *vhp-1*(*sa366*) mutant are not robust to**

**osmotic stress. a)** Trajectory of gonad size vs. body size from hatch to M4 with 95% CI. Graph shows

mean of n = 72 for 0mM NaCl (red) and n = 88 for 250 mM NaCl individuals. **b)** Gonad-to-body ratio

at M4 for same strains as shown in a). The p-values: ANOVA.

### Supplemental Text 1: Derivation of non-dimensionalized resource allocation model

Growth of gonad (g) and soma (s) are both assumed to follow Michaelis-Menten kinetics according to
available nutrient concentration in the respective compartment $N_g/g$ and $N_s/s$ with maximal growth
rate $\mu$ at nutrient saturation:

[1]

$$\frac{dg}{dt} = \dot{g} = \mu \frac{\frac{N_g}{g}}{\frac{N_g}{g} + K} g(t)$$

[2]

$$\frac{ds}{dt} = \dot{s} = \mu \frac{\frac{N_s}{s}}{\frac{N_s}{s} + K} s(t)$$

Total available nutrients are proportional to the size of the soma s and the quality of the environment
$\beta$:

[3]

$$N_{tot}(t) = s(t) * \beta$$

Nutrients N are split between soma and gonad according to the control parameter $\alpha(t)$:

[4]

$$N_g(t) = \alpha(t) * N_{tot}(t) = \alpha(t) * s(t) * \beta$$

[5]

$$N_s(t) = (1 - \alpha(t)) * N_{tot}(t) = (1 - \alpha(t)) * s(t) * \beta$$

Combining eqs. [1,2,4,5] yields:

[6]

$$\frac{\dot{g}(t)}{g(t)} = \mu \frac{\alpha(t) * \beta * s(t) / g(t)}{\alpha(t) * \beta * s(t) / g(t) + K}$$

[7]

$$\frac{\dot{s}(t)}{s(t)} = \mu \frac{(1 - \alpha(t)) * \beta}{(1 - \alpha(t)) * \beta + K}$$

Replacement of time t and environment quality $\beta$ by $\tau = \mu * t$ and $\hat{\beta} = \frac{\beta}{K}$ yields the non-dimensional
model:

[8]

$$\mu_g = \frac{dg/d\tau}{g} = \frac{\dot{g}}{g} = \frac{\alpha(\tau) \hat{\beta} \frac{s(\tau)}{g(\tau)}}{\alpha(\tau) \hat{\beta} \frac{s(\tau)}{g(\tau)} + 1}$$

[9]

$$\mu_s = \frac{ds/d\tau}{s} = \frac{\dot{s}}{s} = \frac{(1 - \alpha(\tau)) \hat{\beta}}{(1 - \alpha(\tau)) \hat{\beta} + 1}$$

### Derivation of special cases of $\alpha(t)$

#### a. Proportional expansion of gonad (g) and soma (s)

Gonad and soma expand proportionally when $\mu_g = \mu_s$ or

[10]

$$\frac{\alpha(\tau) \hat{\beta} \frac{s}{g}}{\alpha(\tau) \hat{\beta} \frac{s}{g} + 1} = \frac{(1 - \alpha(\tau)) \hat{\beta}}{(1 - \alpha(\tau)) \hat{\beta} + 1}$$

Solving eq. 10 for $\alpha(t)$ yields:

[11]
$$\alpha_{phase\ 1}(\tau) = (g(\tau)) / (g(\tau) + s(\tau))$$

**b. Coordinated growth of gonad and soma at constant growth rate ratio $\rho$**

Let $\rho$ be the ratio of the growth rates of gonad and soma

$$[12] \quad \rho = \frac{\mu_g}{\mu_s}$$

Inserting eqs. 8 and 9 yields:

$$[13] \quad \rho = \frac{\alpha(\tau)(1+(1-\alpha(\tau))\hat{\beta})s(\tau)}{(1-\alpha(\tau))(g(\tau)+\alpha(\tau)\hat{\beta}s(\tau))}$$

Solving eq. 13 for $\alpha(\tau)$ yields the positive solution:

$$[14] \quad \alpha_{phase\ 2}(\tau) = \frac{\hat{\beta}\rho s(\tau) - \rho g(\tau) - s(\tau) - \hat{\beta}s(\tau) + \sqrt{4\hat{\beta}(-1+\rho)\rho g(\tau)s(\tau) + (\rho g(\tau) + (1+\hat{\beta} - \hat{\beta}\rho)s(\tau))^2}}{2\hat{\beta}(\rho-1)s(\tau)}$$

with $\rho$ to reach a given adult gonad-to-soma proportion being

$$[15] \quad \rho = \frac{\text{Log}((s_{adult}/(g_{adult}/s_{adult}))/g_{start})}{(1-\alpha)(g(\tau)+\alpha\hat{\beta}s(\tau))}$$

**c. Bang-bang control**

Bang-bang control was implemented by $\alpha(\tau) = 0$ until $s = s_{adult}$, such that:

$$[16] \quad \frac{\dot{s}}{s(\tau)} = \frac{\hat{\beta}s(\tau)}{1+\hat{\beta}}, \frac{\dot{g}}{g(\tau)} = 0$$

followed by $\alpha(\tau) = 1$ until $g = g_{adult}$, such that

$$[17] \quad \frac{\dot{s}}{s(\tau)} = 0, \frac{\dot{g}}{g(\tau)} = \frac{\hat{\beta}s_{adult}}{g(\tau)+\hat{\beta}s_{adult}}$$

For the equations of the nondimensionalized model shown in the main text, the hat on $\beta$ was omitted for simplicity, and variable $t$ instead of $\tau$ is used for nondimensionalised time.
